# Antibiotic Resistance pattern in Bile from Cholecystectomised Patients by Multiplex PCR

**DOI:** 10.1101/2024.06.25.600685

**Authors:** Nasreen Farhana, Mohammad Ashraf Uddin Khan, SM Shamsuzzaman

## Abstract

**Background:** Bile in the biliary tract is normally sterile. Presence of gallstones, ascending infection from duodenum or bacterial translocation from portal vein leads to microfloral colonization in biliary system.

**Aim:** This study was aimed to evaluate the microbiological profile of bile from gall bladder for determination of the appropriate antibiotics in cholecystectomised patients.

**Methods:** This cross sectional study included patients who underwent laparoscopic or open cholecystectomy in Dhaka Medical College Hospital, Dhaka, from July, 2013 to December, 2014. Total 246 intraoperative bile were cultured aerobically in Blood agar and MacConkey’s agar media. The identified isolates were tested for their sensitivity pattern according to CLSI guidelines and multiplex PCR was used to detect virulence genes of *Salmonella* Typhi and anaerobic bacteria along with drug resistance genes.

**Findings:** Out of 246 bile samples, organisms were identified in 171 (69.51%) cases; 119 (48.37%) were aerobic bacteria identified by culture and PCR and 52 (21.14%) were anaerobic bacteria identified by multiplex PCR. *Escherichia coli* (26.61%) were found predominantly followed by *Staphylococcus aureus* (19.35%), *Clostridium perfringens* (13.82%). *Salmonella enterica* serovar Typhi was detected by culture and PCR in 3 (1.22%) and 8 (3.45%) samples respectively. Prevalence of ESBLs, Carbapenemase producers and MRSA were detected phenotypically in 10.96%, 16.44% and 8.33% samples respectively and the resistance genes blaCTX-M-15 (50.0%), blaOXA-1-group (25.0%), blaNDM-1 (62.50%), OXA-181/OXA-84 (12.5%) and mecA (8.33%) were detected.

**Conclusion:** Significant proportion of aerobic and anaerobic bacterial infection associated with biliary tract obstruction may warrants serious health risk to cholecystectomised patients in this region.

## Introduction

Bile in the gallbladder is usually sterile. Biliary infection is a result of biliary obstruction due to symptomatic cholelithiasis, common bile duct stones, cholecystitis or cholangitis [1] and indicate the presence of microflora in 20% - 46% of the patients.[2-4] Aerobic bacterial infection is the most common with gram negative preponderance while viral, fungal agents and anaerobic bacteria are uncommon causative agents.[5]

Monomicrobial growths are more frequent (30.8% - 96%) in comparison with polymicrobial cultures (4% - 4.8%).[6-8] The commonly isolated aerobic bacteria from culture of bile are *Escherichia coli* (17.5% - 62%)[2,9] *Pseudomonas* spp. (5.4% - 23.7%),[7,10] *Acinetobacter* spp. (2.7% - 5.63%),[6,11] *Klebsiella* spp. (5.3% -27%),[4,8] *Staphylococcus aureus* (2.7% - 7.04%),[6,11] *Enteroccocus* spp. (13.1% -15.6%),[4,12] *Citrobacter* spp. (5.63% - 9.5%),[10,11] *Salmonella enterica* serovar Typhi (2.82% -14%).[11,13] β-glucuronidase producing anaerobes especially *Bacteroides fragilis* (12.86%-58.82%) and *Clostridium perfringens* (32.86% - 36.58%) are recovered in both acutely inflamed and chronically inflamed biliary systems.[14,15]

Survival of *Salmonella enterica* serovar Typhi in the gallbladder seems to involve several strategies - invasion of the gallbladder epithelium, escaping from the extremely high concentrations of bile salts and formation of bile inducing biofilms on gallstones in gallbladder lumen.[16,17] In general, 2-5% of all individuals who develop clinical or subclinical infection with *S*. Typhi become chronic gallbladder carriers forming biofilms which possess the Vi antigen associated with virulence and continue to intermittently shed organisms for a prolonged period which are generally more numerous in the bile than feces.[18-20]

The most effective treatment for S. Typhi carrier state is a combination of surgery and antibiotics. Antimicrobial agents administered empirically should be changed according to their sensitivity suggesting of recent changes in antibiotic-resistant profiles in the patients.[21] The major problem for infection control is the spread of extended spectrum β-lactamase (ESBL)- producing organisms which confers resistance to most β-lactam antibiotics.[22] In addition, the emergence of carbapenemase-producing bacteria has a significant impact on clinical outcomes and risk factors associated with prior biliary intervention, nosocomial infection, recurrent cholangitis and indwelling biliary devices.[22,23]

Data regarding bacteriological profile from bile is scanty in Bangladesh. This study was designed to isolate aerobic bacteria by culture and to see their antibiotic susceptibility pattern and detect drug resistance and virulence genes of the anaerobic bacteria.

## Materials and Methods

This cross sectional study was conducted in the Department of Microbiology, Dhaka Medical College from July 2013 to December 2014 after approval from the research review committee (RRC) and ethical review committee (ERC) of Dhaka Medical College. Total 246 patients underwent open or laparoscopic cholecystectomy irrespective of age and sex were included in this study after obtaining informed written consents from the patients according to the Helsinki Declaration. Patient’s age, sex, clinical history, antibiotic history, ultrasonography reports and information about cholecystectomy operation were recorded. Non-probability consecutive sampling technique were used and the sample size was calculated by using formula with 20% prevalence of bacteria isolated from the bile at 95% level of confidence with 5% margin of error. [18] The statistical significance was assigned a p value of <0.05 using the z-test of proportion.

About 3 ml of bile was aspirated from gallbladder during cholecystectomy and carried to the microbiology laboratory of Dhaka Medical College in an hour.[4] The aspirated bile was immediately inoculated on to 5% sheep blood agar, chocolate agar and MacConkey agar media and incubated aerobically at 37°C and examined after 24-48 hours. All bacterial isolated were identified by colony morphology, gram staining and relevant biochemical tests.[24,25]

### Antimicrobial susceptibility test

Antimicrobial susceptibility of the isolated bacteria was done by Kirby-Bauer disc diffusion technique using commercially available antibiotic disks (Oxoid Ltd, Basingstoke, United Kingdom) according to Clinical and Laboratory Standard Institute (CLSI) guidelines.[26,27]

### Isolation of drug resistant bacteria

All *S. aureus* isolates were screened by standard disc diffusion method and reported as MRSA as per CLSI guidelines.[27] All MRSA isolates were compared with MIC of oxacillin and confirmed by detection of mecA and PVL gene by PCR using specific primers (Table 1).[28,29] ESBLs and carbapenemase producing gram negative bacilli were phenotypically detected by double disc synergy (DDS) test, combined disc (CD) assay and modified Hodge Test (MHT).[30-32]

**Table 1:**
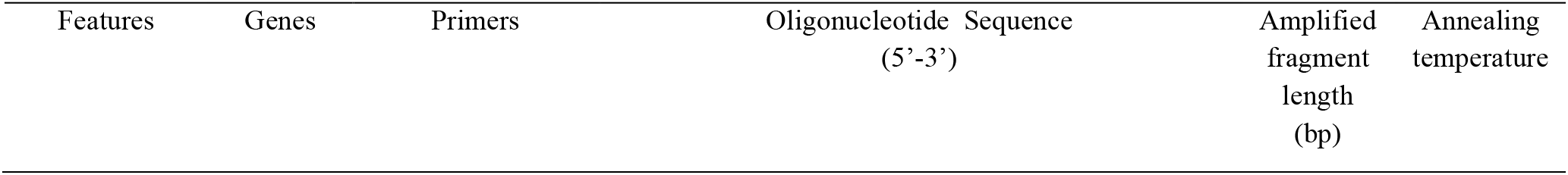
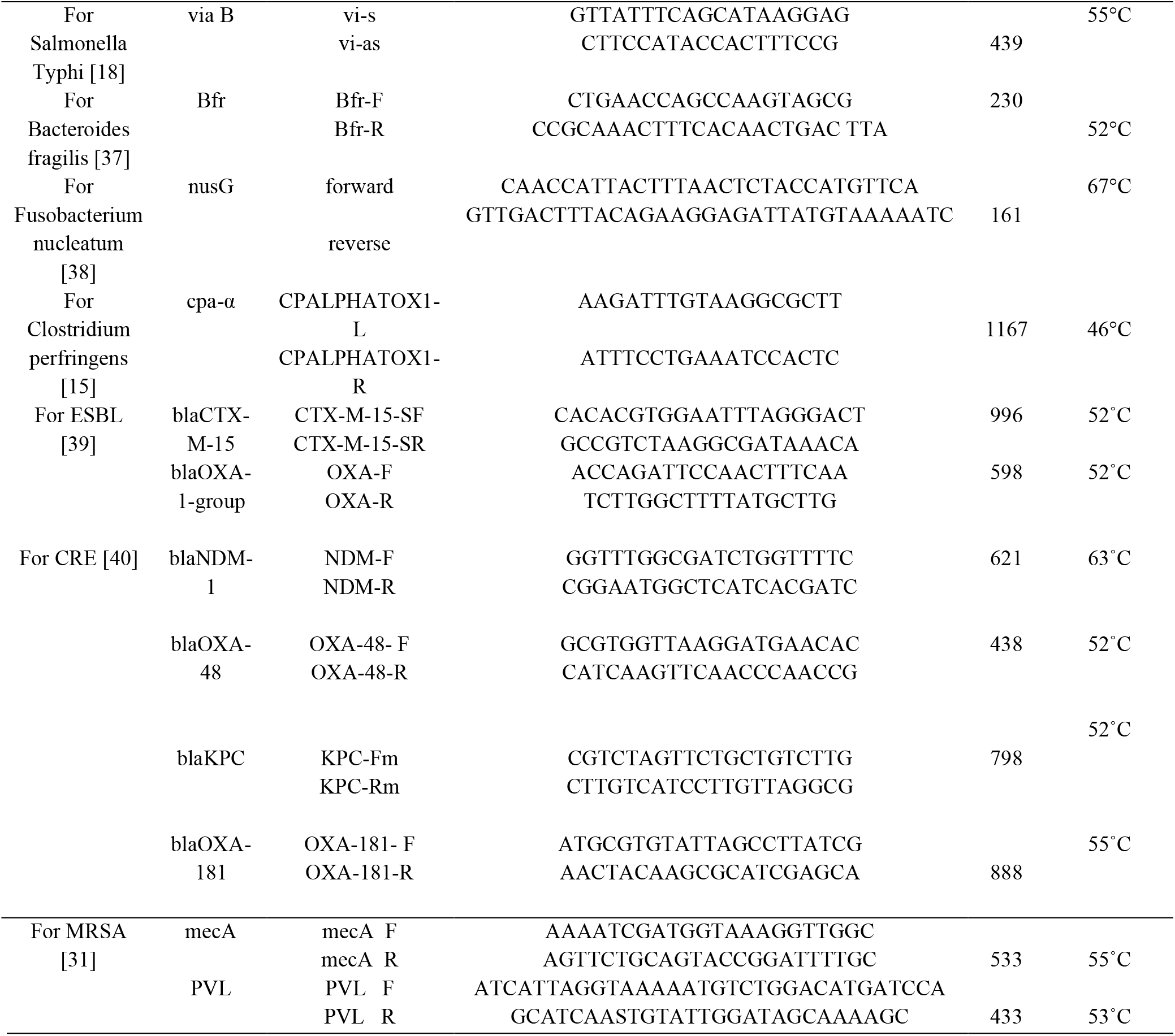
The primers were used in this study.

### Extraction and amplification of virulence genes

Total DNA was extracted from each bile sample by boiling method. About three milliliters of bile were pelleted by centrifugation for 5 min at 14,000g, incubated with 100 µl lytic buffer [composition: 50mM Tris-HCl (pH 8.0), proteinase K (50µg/ml) and 0.5% tween 20 solution] (Promega corporation, USA) for 2 hour at 55°C after vortexing thoroughly and placed in a heat block (DAIHA Scientific, Seoul, Korea) at 100°C for 10 minutes for boiling. After immediately transferred to the ice the mixtures kept for 5 minutes and centrifuged at 10,000 g at 4°C for 15 minutes. The supernatant were used as template DNA and preserved at 4⁰C for 7-10 days and -20°C for long time. [33] Specific primers were mixed with Tris-EDTA (TE) buffer according to manufacturer’s instruction (Table 1). PCR was performed in a thermal cycler (Eppendorf AG, Master cycler gradient, Hamburg, Germany) with final reaction volume of 25µl containing 10 µl of mastermix [premixed mixture of 50 units/ml of Taq DNA polymerase supplied in a proprietary reaction buffer (pH 8.5), 400μM dNTP, 3mM MgCl2 (Promega corporation, USA)], 1 µl forward primer and 1 µl reverse primer (Promega corporation, USA), 3 µl extracted DNA and 14 µl nuclease free water and initial denaturation at 95°C for 10 minutes followed by 35 cycles including annealing temperature which varied in different reaction, elongation at 72°C for 1 min and a final extension at 72°C for 10 minutes. (Table 1) [15,18,34-36]

### Analysis of PCR products

The PCR products were electrophoresed in 1.5% agarose gel, staining with ethidium bromide destaining with distilled water and observed under UV Transilluminator (Gel Doc, Major science, Taiwan) for DNA bands which were identified according to their molecular size by comparing with 100 bp DNA ladder. [30]

## Results

In this study 246 cholecystectomised patients were between the ages of 20 to 70 years and 73 (29.68%) of them were between 40-50 years of age. Male patients were 64 (26.02%) and female were 182 (73.98%) and male female ratio was 1:2.84 (Table 2).

**Table 2:** Sex distribution in relation to organisms detected (n=246).

| Sex | No growth on culture | Detection of positive cases for organism |  | Total positive cases n (%) | p value |
| --- | --- | --- | --- | --- | --- |
|  |  | Aerobic bacteria by culture and PCR | Anaerobic bacteria by PCR* |  |  |
|  | n (%) | n (%) | n (%) |  |  |
| Male (n=64) | 34 (53.13) | 31a (48.44) | 15 (23.44) | 46 (71.88) | p > 0.05 |
| Female (n=182) | 98 (53.85) | 88b (48.35) | 37 (20.33) | 125 (68.68) |  |
| Total (n=246) | 132 (53.66) | 119 (48.37) | 52 (21.14) | 171 (69.51) |  |

Note: Male female ratio = 1:2.84
‘\*’= Anaerobic culture was not done.
‘a’= Out of 31 aerobic bacteria, one was positive for S. Typhi by PCR.
‘b’ = Out of 88 aerobic bacteria, 4 were positive for S. Typhi by PCR.

Out of 246 aspirated bile samples, there was no growth on cultures in 132 (53.66%) cases while organisms were identified in 171 (69.51%) cases by both culture and multiplex PCR (Table 2). Out of 64 male and 182 female patients, organisms were identified in 46 (71.88%) and in 125 (68.68%) cases respectively (p > 0.05).

Aerobic bacteria were identified in 119 (48.37%) cases by both culture and PCR and of them 7 (13.46%) were mixed with anaerobic bacteria (Table 2). Anaerobic bacteria were identified by PCR in 52 (21.14%) cases and *Clostridium perfringens, Fusobacterium nucleatum* and *Bacteroides fragilis* were in 34 (13.82%), 10 (4.07%) and 8 (3.25%) cases respectively (Figure 1, 2).

**Figure 1:**
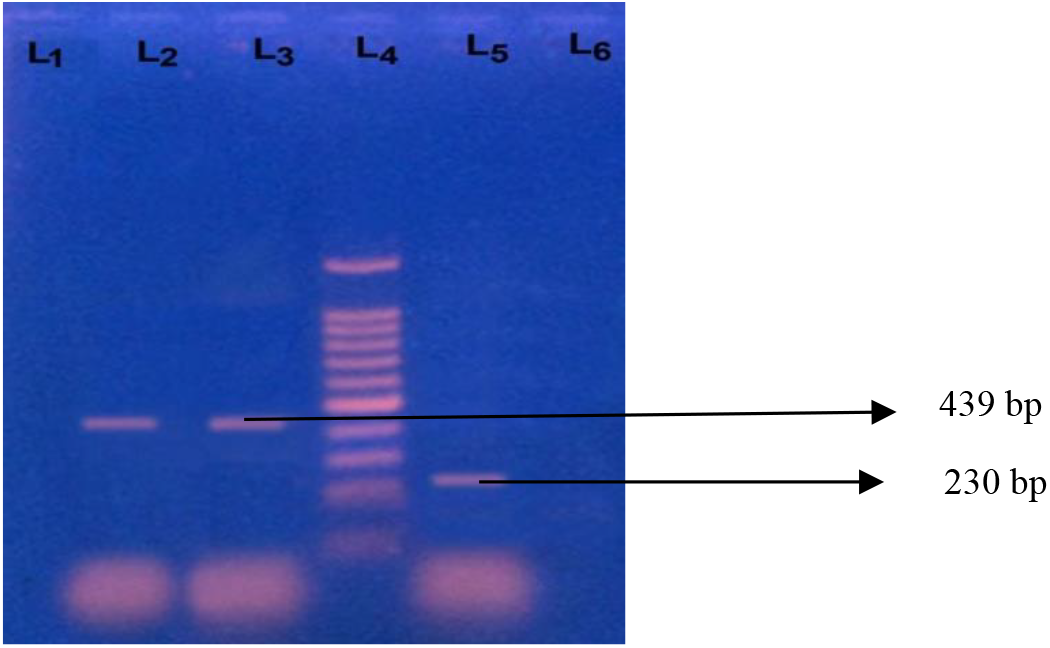
Photograph of amplified DNA of Salmonella Typhi and Bacteroides fragilis. Lane 1: Negative control (DNA of Pseudomonas aeruginosa). Lane 2: Positive control of Salmonella Typhi. Lane 3: Bile sample (positive for Salmonella Typhi of 439 bp). Lane 4: 100bp DNA ladder. Lane 5: Bile sample (positive for Bacteroides fragilis of 230 bp).

**Figure 2:**
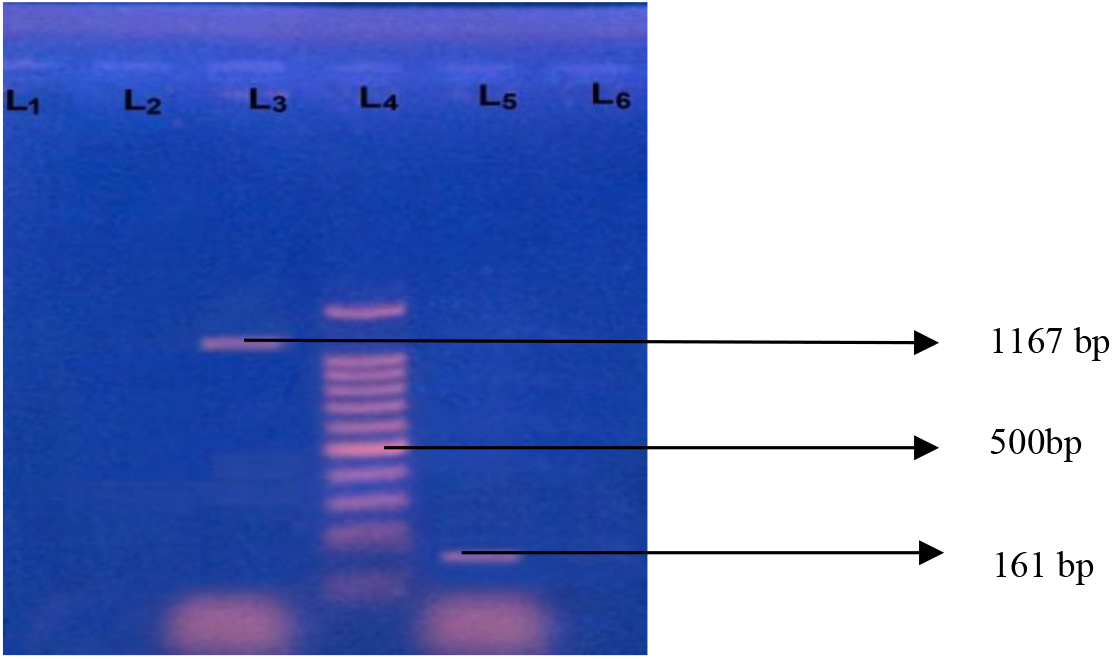
Photograph of amplified DNA of Clostridium perfringens and Fusobacterium nucleatum. Lane 1: Negative control (DNA of Fusobacterium necrophorum). Lane 2: Bile sample (positive for Clostridium perfringens of 1167 bp). Lane 4: 100bp DNA ladder. Lane 5: Bile sample (positive for Fusobacterium nucleatum of 161 bp).

Among the 119 aerobic bacteria, 114 (46.34%) were culture positive and of them 83 (66.94%) were gram negative bacilli and 41 (33.06%) were gram positive cocci. Among them, *Escherichia coli* (26.61%), *Staphylococcus aureus* (19.35%), *Citrobacter freundii* (14.52%), *Pseudomonas* spp. (11.29%), *Viridans streptococcus* (8.06%), *Acinetobacter* spp. (3.51%), *Salmonella enterica* serovar Typhi (2.63%) and *Klebsiella* spp. (1.75%) were predominantly found (Table 3). *Salmonella* Typhi 5 (4.20%) was also identified by PCR (Figure 1).

**Table 3:** Aerobic organisms isolated from bile of study population.

| Organism | Culture positive |  | Total<br>n (%) |
| --- | --- | --- | --- |
|  | Monomicrobial<br>n (%) | Polymicrobial<br>n (%) |  |
| Escherichia coli | 30 (90.91) | 3 (9.09) | 33 (26.61) |
| Pseudomonas spp. | 10 (71.43) | 4 (28.57) | 14 (11.29) |
| Klebsiella pneumoniae | 2 (100.00) | 0 (0.00) | 2 (1.61) |
| Klebsiella oxytoca | 2 (100.00) | 0 (0.00) | 2 (1.61) |
| Citrobacter freundii | 14 (77.77) | 4 (22.22) | 18 (14.52) |
| Citrobacter koseri | 5 (71.43) | 2 (28.57) | 7 (5.65) |
| Acinetobacter spp. | 4 (100.00) | 0 (0.00) | 4 (3.23) |
| Salmonella enterica serovar Typhi | 1 (33.33) | 2 (66.67) | 3 (2.42) |
| Staphylococcus aureus | 21 (87.50) | 3 (12.50) | 24 (19.35) |
| CONS | 5 (100.00) | 0 (0.00) | 5 (4.03) |
| Viridans streptococci | 9 (90.00) | 1 (10.00) | 10 (8.06) |
| Streptococcus pneumoniae | 2 (100.00) | 0 (0.00) | 2 (1.61) |
| Total | 105 (84.68) | 19 (15.32) | 124 (100.00) |

\* CONS = Coagulase negative Staphylococcus

**Table 4:** Antibiotic resistance pattern of isolated organisms to different antibiotics.

| Antimicrobial drugs | Escherichia coli (n=30) | Pseudomonas spp. (n=12) | Citrobacter freundii (n=14) | Citrobacter koseri (n=6) | Acinetobacter spp. (n=4) | Klebsiella pneumoniae | Klebsiella oxytoca (n=2) | Salmonella serovar Typhi (n=03) | Staphylococcus aureus | Coagulase negative staphylococcus | Viridans Streptococcus (n (%)) | Streptococcus pneum |
| --- | --- | --- | --- | --- | --- | --- | --- | --- | --- | --- | --- | --- |

|  | n (%) | n (%) | n (%) | n (%) | n (%) | (n=2)<br>n (%) | n (%) | n (%) | n (%) | occus<br>n (%) |  | oniae<br>n (%) |
| --- | --- | --- | --- | --- | --- | --- | --- | --- | --- | --- | --- | --- |
| Ceftriaxone | 22<br>(73.33) | 5<br>(41.67) | 9<br>(64.29) | 4<br>(66.67) | 0 (0.00) | 1<br>(50.00) | 2<br>(100.00) | 0 (0.00) | 9<br>(37.50) | 1 (20.00) | 2 (20.00) | 0<br>(0.00) |
| Amoxiclav | 20<br>(66.67) | 7<br>(58.33) | 9<br>(64.29) | 3<br>(50.00) | 0 (0.00) | 2<br>(100.00) | 2<br>(100.00) | 1 (33.33) | 0 (0.00) | 0 (0.00) | 0 (0.00) | 0<br>(0.00) |
| Ceftazidime | 18<br>(60.00) | 7<br>(58.33) | 9<br>(64.29) | 3<br>(50.00) | 2<br>(50.00) | 2<br>(100.00) | 2<br>(100.00) | 0 (0.00) | - | - | - | - |
| Cefotaxime | 24<br>(80.00) | 6<br>(50.00) | 5<br>(35.71) | 2<br>(33.33) | 1<br>(25.00) | 0 (0.00) | 2<br>(100.00) | 0 (0.00) | - | - | - | - |
| Cefexime | 12<br>(40.00) | 2<br>(16.67) | 2<br>(14.29) | 2<br>(33.33) | 0 (0.00) | 0 (0.00) | 1 (50.00) | 1 (33.33) | - | - | - | - |
| Ciprofloxacin | 12<br>(40.00) | 1 (8.33) | 4<br>(28.57) | 1<br>(16.67) | 0 (0.00) | 0 (0.00) | 0 (0.00) | 0 (0.00) | 0 (0.00) | 0 (0.00) | 0 (0.00) | 0<br>(0.00) |
| Aztreonam | 6<br>(20.00) | 2<br>(16.67) | 1 (7.14) | 1<br>(16.67) | 1<br>(25.00) | 2<br>(100.00) | 0 (0.00) | - |  |  |  |  |
| Amikacin | 8<br>(26.67) | 1(8.33) | 5<br>(35.71) | 0(0.00) | 0 (0.00) | 0 (0.00) | 0 (0.00) | 1 (33.33) | 0 (0.00) | 0 (0.00) | 2 (20.00) | 1<br>(50.00) |
| Gentamicin | 8<br>(26.67) | 4<br>(33.33) | 4<br>(28.57) | 0<br>(0.00) | 0 (0.00) | 0 (0.00) | 0 (0.00) | 0 (0.00) | 0 (0.00) | 0 (0.00) | 0 (0.00) | 0<br>(0.00) |
| Meropenem | 2 (6.67) | 0 (0.00) | 0 (0.00) | 0<br>(0.00) | 0 (0.00) | 0 (0.00) | 0 (0.00) | 0 (0.00) | - | - | - | - |
| Imipenem | 6<br>(20.00) | 0 (0.00) | 1 (7.14) | 0<br>(0.00) | 0 (0.00) | 1<br>(50.00) | 0 (0.00) | 0 (0.00) | - | - | - | - |
| Colistin | 4<br>(13.33) | 1 (8.33) | 0 (0.00) | 0<br>(0.00) | 0 (0.00) | 0 (0.00) | 0 (0.00) | 0 (0.00) | - | - | - | - |
| Piperacillin -<br>Tazobactam | 4<br>(13.33) | 0 (0.00) | 0 (0.00) | 0<br>(0.00) | 0 (0.00) | 0 (0.00) | 0 (0.00) | - | - | - | - | - |
| Azithromycin | 2 (6.67) | 2<br>(16.67) | 5<br>(35.71) | 1<br>(16.67) | 1<br>(25.00) | 1<br>(50.00) | 1 (50.00) | 1 (33.33) | 0 (0.00) | 1 (20.00) | 2 (20.00) | 1<br>(50.00) |
| Nalidixic acid | - | - | - | - | - | - | - | 1 (33.33) | - | - | - | - |
| Novobiocin | - | - | - | - | - | - | - | - | 0 (0.00) | 0 (0.00) | - | - |
| Cefoxitin | - | - | - | - | - | - | - | - | 2 (8.33) | 0 (0.00) | - | - |
| Oxacillin | - | - | - | - | - | - | - | - | 2 (8.33) | 0 (0.00) | - | - |
| Vancomycin | - | - | - | - | - | - | - | - | 0 (0.00) | 0 (0.00) | 0 (0.00) | 0<br>(0.00) |
| Linezolid | - | - | - | - | - | - | - | - | 0 (0.00) | 0 (0.00) | 0 (0.00) | 0<br>(0.00) |
| Clindamycin | - | - | - | - | - | - | - | - | 0 (0.00) | 0 (0.00) | 1 (10.00) | 0<br>(0.00) |
| Optochin | - | - | - | - | - | - | - | - | - | - | 10<br>(100.00) | 0<br>(0.00) |
| Doxycycline | - | - | - | - | - | - | - | - | 4<br>(16.67) | 1 (20.00) | 1 (10.00) | 0<br>(0.00) |
| Levofloxacin | - | - | - | - | - | - | - | - | 0 (0.00) | 0 (0.00) | 0 (0.00) | 0<br>(0.00) |

Note: (-) = not used.

Single organisms were predominantly isolated in 105 (84.68%) cases and polymicrobial infection was detected in 9 (15.32%) cases and among them, 8 (7.02%) had 2 organisms and one (0.87%) had 3 organisms. There was no definite predominance of any combination in polymicrobial infection (Table 3).

Most of the isolated organisms were resistant to third generation cephalosporins but susceptible to carbapenems, piperacillin-tazobactam and colistin. *Salmonella* Typhi was found most sensitive to ceftriaxone and ciprofloxacin (Table 5). ESBL and carbapenemase producing gram negative bacteria and MRSA were found in 16.44%, 10.96% and 8.33% cases respectively.

The amplified ESBL encoding genes using multiplex PCR revealed fragments for *bla*CTX-M-15 and *bla*OXA-1-group and were positive in 50% and 25% of ESBL producers respectively. Two organisms had both *bla*CTX-M-15 and *bla*OXA-1-group of genes (Figure 3).

**Fig 3:**
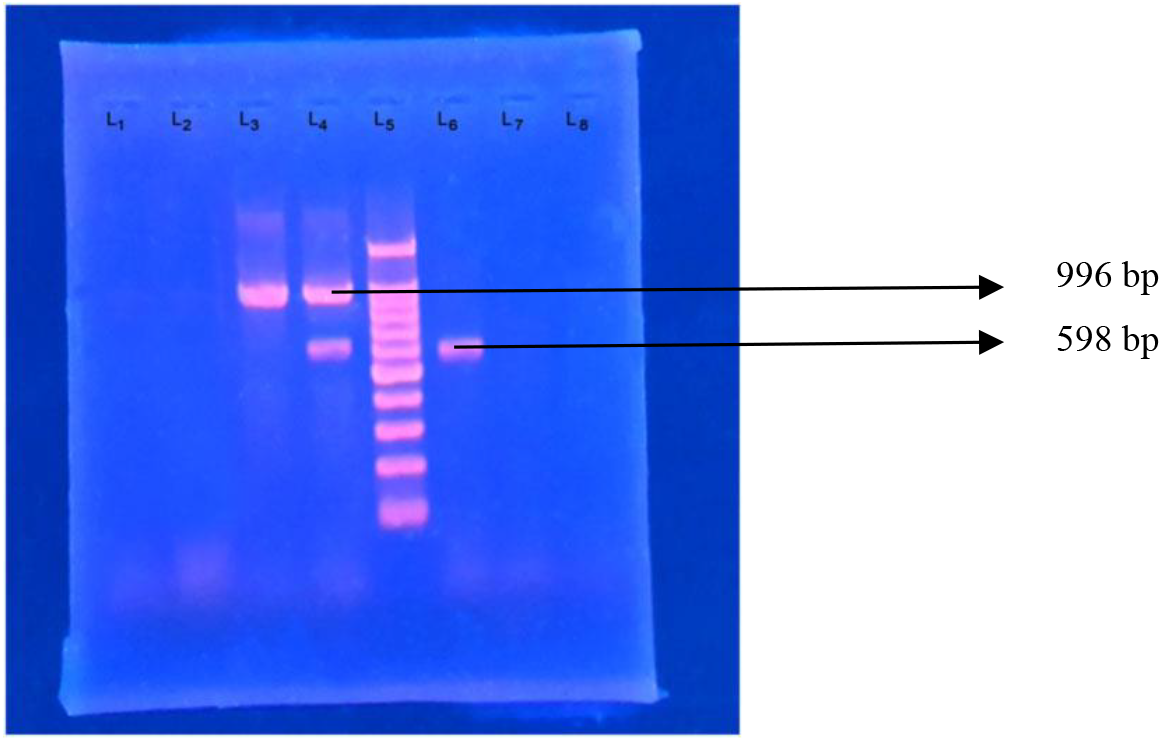
Photograph of amplified DNA of ESBL encoding genes. Lane 1: Negative control without DNA, Lane 3: amplified DNA of 996 bp for blaCTX-M-15 gene, Lane 4: amplified DNA of 598 bp for blaOXA-1 group gene and 996 bp for blaCTX-M-15 gene, Lane 5: 100 bp DNA ladder and Lane 6 detecting amplified DNA of 598 bp for blaOXA-1 group gene.

Carbapenemase encoding genes NDM-1 producers were the most prevalent (62.5%) whilst OXA-181/OXA-84 was the least common carbapenemase (12.5%) and none was found KPC producer (Figure 4). All the 2 MRSA strains were mecA positive and none was PVL gene positive (Figure 5).

**Figure 4:**
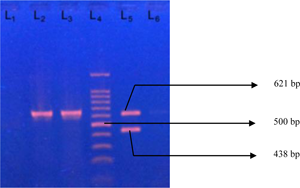
Photograph of amplified DNA of carbapenemase encoding genes. Lane 2: positive control of blaNDM-1, Lane 3: amplified blaNDM-1 (621 bp), Lane 4: 100 bp DNA ladder, Lane 5: both blaNDM-1 (621 bp) and blaOXA-48 (438 bp) and Lane 7: negative control– Escherichia coli ATCC 25922.

**Figure 5:**
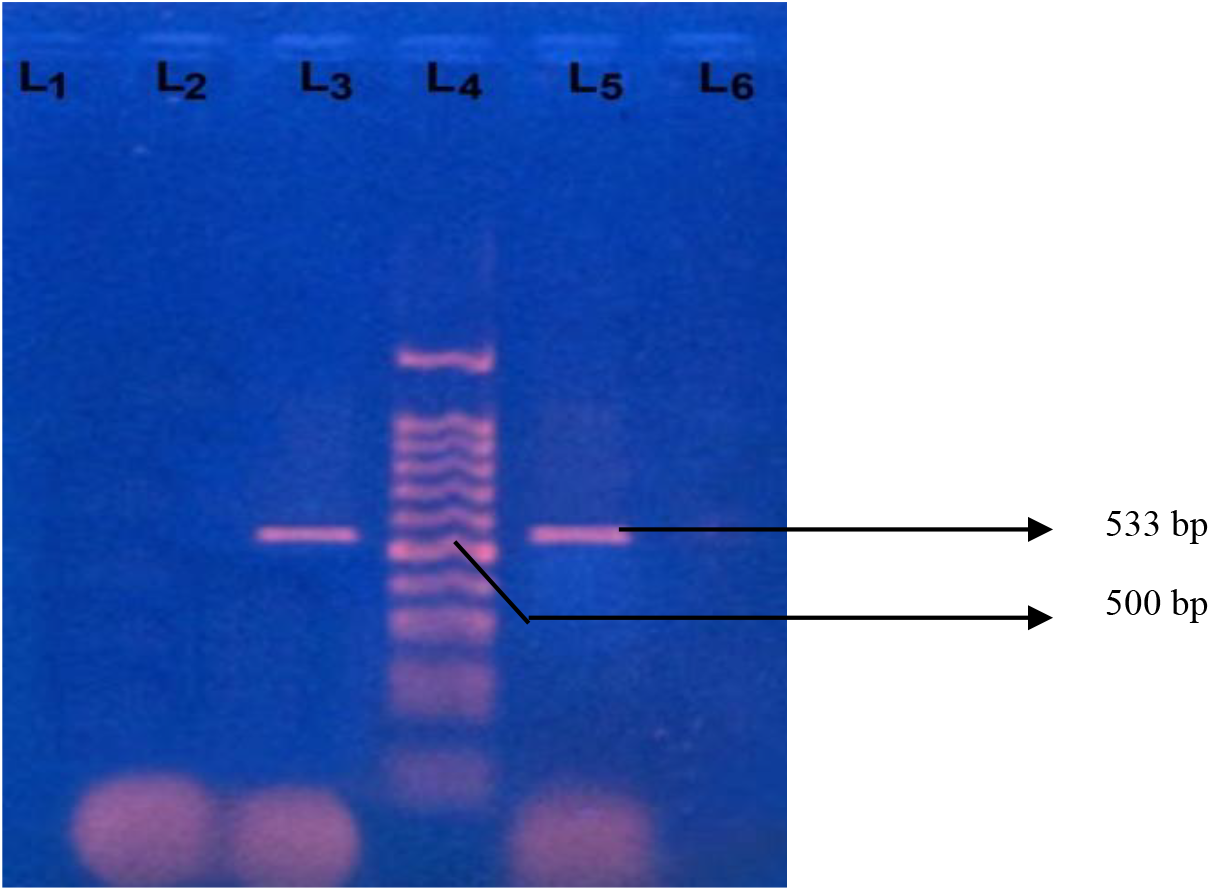
Photograph of amplified DNA of mecA gene (533 bp) in lane 3 and 5. Lane 2: negative control - Staphylococcus aureus ATCC 25923 and Lane 4: 100 base pair DNA ladder.

## Discussion

In consistent with the present study, a number of studies revealed that incidence of biliary obstruction due to gallstones is common above the age of forty and the proportions were 25-53%.[8,37] Predominance of female patients were found in this study and attributed to estrogens with HMG-Co-A reductase enzyme causing increased formation of cholesterol stones.[37]

In the present study aerobic bacteria were identified by both culture and PCR in 119 (48.37%) cases and anaerobic bacteria were identified by multiplex PCR in 52 (21.14%) cases. In contrast to this study, Ballal et al. had detected higher proportion of aerobic (56.8%) and lower proportion of anaerobic (13.6%) bacteria from 50 patients with biliary tract disease.[11] The reason behind the lower proportion of aerobic bacteria from bile in the present study might be due to the fact that many patients might have taken antibiotics before cholecystectomy. Mixed aerobes and anaerobes were identified in this study by both culture and multiplex PCR in 7 (13.46%) cases while some studies found in 5-6% cases from bile cultures.[38,39] Among anaerobic bacteria β-glucuronidase producing *Cl. perfringens* and *B. fragilis* were the most frequent isolates which was also supported by Atia et al.[14] β-glucuronidase catalyze the hydrolysis of bilirubin glucuronide to form insoluble salts that plays a role in gallstone formation.[15]

The present study showed positive bile culture in 114 (46.34%) patients which was comparable to other study with isolation rates of 24-50.5%.[4,7] In a study by Dongol et al. showed a lower proportion (20%, 274/1377) of positive bile cultures with only gram negative bacteria while Ahmad et al. has reported considerably higher (58.58%, 157/268) culture positive results than the present study.[8,18] Polymicrobial infection has been reported in 4-7% of bile aspirates while in the present study, polymicrobial growth was detected in 9 (15.32%) cases.[7,40] The increased rate of polymicrobial growth may be due to some antibiotics resistance bacteria along with food habits and environmental factors. The most common gram negative bacilli isolated in the present study were *Esch. coli* (26.61%) followed by *C. freundii* (14.52%) and *Pseudomonas* spp. (11.29%). *Esch. coli* has been also reported as the predominant flora by other studies [8,13,18] which attributed to the activity of its β glucuronidase enzyme in formation of calcium billirubinate gallstone formation.[13,39] This is in contrast to the study by Gill et al. where the most common organisms isolated were *Pseudomonas* spp. (68.42%).[41] Out of 246 bile samples, *S*. Typhi was detected in 3 (1.22%) by culture and in 8 (3.25%) by PCR which reported similar in different studies [18,41] but higher prevalence (8-14%) were reported by some studies which might be due to hyperendemicity of the study area.[8,13,41] Of gram-positive cocci, *S. aureus* was found in 24 (19.35%) cases in this study which correlates with the study by Suri et al.[42] Lower isolation rate (2.1%) and higher isolation rate (29.67%) were reported in different studies which might be due to geographical variation and antibiotic policy of the respective country.[7,39]

High level of resistance to third generation cephalosporins was observed in some studies which is similar to the present study.[8,36] The most high resistance profiles in this study were *E. coli, P. aeruginosa, Klebsiella* spp. and *C. freundii* similar to the study by Kaya et al.[7] *S*. Typhi was found most susceptible to ciprofloxacin and ceftriaxone while another study by Dongol et al. had showed resistance to only nalidixic acid in *Salmonella* from bile.[18] The isolated gram positive bacteria were found resistant to ceftriaxone, doxycycline, oxacillin and cefoxitin which was supported by Ahmad et al.[8] Among the isolated MRSA was found sensitive to vancomycin and linezolid which coincides with a study by Shahi et al.[40]

ESBLs producing gram negative bacilli are increasingly prevalent from patients with gallstone disease whereas a high presence of CTX-M-type ESBL genes was identified using real-time PCR and pyrosequencing by a number of studies which differ from the present study.[35,38,39] The prevalence of *bla*NDM-1 gene in gram negative bacilli in bile samples is due to healthcare associated acquisition in hospitalized patients.[36,43] Therefore the evaluation of antibiotic susceptibility patterns in isolated bacteria and screening of drug resistance genes in bile samples are essential in hospitalized patients.

## Conclusion

Significant proportions of both aerobic and anaerobic bacteria were detected from bile samples by culture and PCR. Meropenem, piperacillin-tazobactam, colistin and vancomycin may be a part of empirical regime associated with antibiotic resistant bacteria in cholecystectomised patients and further study on antibiotic susceptibility in those patients is warranted.

## Acknowledgments

We acknowledged the support of the cholecystectomised patients and the Department of Microbiology, Dhaka Medical College, Dhaka for providing the opportunity and resources to undertake the study.

